# Controlling reactogenicity while preserving immunogenicity from a self-amplifying RNA vaccine by modulating nucleocytoplasmic transport

**DOI:** 10.1101/2024.12.29.630334

**Authors:** Jason A Wojcechowskyj, Robyn M Jong, Imre Mäger, Britta Flach, Paul V Munson, Pragya Mukherjee, Barbara Mertins, Katherine R Barcay, Thomas Folliard

## Abstract

Self-amplifying RNA (saRNA)-based vaccines have emerged as a potent and durable RNA vaccine platform. However, RNA vaccine platforms trigger undesirable side effects at protective doses, underscoring the need for improved tolerability. To address this, we leveraged the Cardiovirus leader protein, which is well-characterized to dampen host innate signaling by modulating nucleocytoplasmic transport (NCT). Co-administration of a leader-protein-encoding mRNA (which we have named “RNAx”) delivered alongside vaccine cargo saRNA reduced interferon production while enhancing Influenza hemagglutinin (HA) expression in human primary cells and murine models. RNAx potently decreased serum biomarkers of reactogenicity after immunizations with an HA-expressing saRNA-LNP vaccine while maintaining the magnitude of the antibody and cellular response. RNAx also consistently enhanced binding antibody titers after a single injection and in some conditions enhanced binding antibody and neutralization titers post-boost. These findings support RNAx as a promising platform approach for improving tolerability of saRNA-LNP vaccines while preserving or enhancing immunogenicity.

## INTRODUCTION

RNA lipid nanoparticle (LNP) vaccines represent a breakthrough in vaccinology, offering unprecedented speed in responding to emerging pathogens while providing robust protection and simplified manufacturing^1^. However, despite the public health success of SARS-CoV-2 mRNA-LNP vaccines, these vaccines exhibit undesirable side effects at protective µg doses levels^2^ and are expensive to manufacture rapidly on a global scale^3^. Self-amplifying RNA (saRNA) vaccines are compelling alternatives to conventional mRNA platforms, providing longer lasting protection at lower µg doses^4,5^. The higher potency per µg dose from saRNA is likely a product of the combination of robust antigen expression in the presence of higher levels of adjuvancy arising from innate signaling, inherent to the saRNA platform. Self-replication in a cell generates double-stranded RNA (dsRNA) intermediates, a potent adjuvant that itself can contribute to systemic reactogenicity^6–8^. While innate pathways provide crucial adjuvant effects, their overactivation can trigger reactogenicity and suppress immune responses from RNA-based vaccines, for example, by inhibiting antigen expression. saRNA and dsRNA are potent inducers of primary interferons (IFN-α/β), which have been shown to inhibit both T cell^9,10^ and B cell responses^11–14^ from RNA vaccines. Significant research has been made to balance saRNA-induced innate signaling via co-delivering small molecules^11,15^ or encoded protein-based^11,16^ inhibitors of innate signaling, incorporating modified nucleosides^17,18^, mutating the saRNA vector^19^, or employing non-LNP-based delivery formulations^12,20,21^. While these approaches show considerable promise, there remains an opportunity to develop strategies that combine low µg doses with effective suppression of reactogenicity.

Here we report the engineering of the leader protein from Cardiovirus, a genus of RNA viruses from the Picornaviridae family that broadly dampens excessive innate immune activation triggered by saRNA. The leader protein inhibits innate signaling and interferon (IFN) production^22^ driven by the IRF-3 transcription factor^23^ and disrupts translational inhibition induced by PKR activation^24^ through interference with nucleocytoplasmic transport (NCT)^25–28^. This mechanism of suppressing host gene expression is essential for viral gene expression and replication in IFN-competent hosts^29,30^. We leveraged this function of the leader protein to balance excessive saRNA-driven innate signaling and IFN production in the context of an saRNA vaccine. Here, we encode the leader protein from a small mRNA termed “RNAx”. A key advantage of RNAx is that it targets NCT broadly, thus enabling suppression of diverse innate signaling pathways simultaneously^31^.

From experiments in cell lines, primary human immune cells, and mouse models, we find that RNAx suppresses saRNA-induced IFN production and other proinflammatory cytokines while enhancing antigen expression both *in vitro* and *in vivo*. We also find that RNAx enhances gene of interest (GOI) expression and suppresses proinflammatory cytokines from unmodified mRNA-LNPs in mice. Following administration of an saRNA-LNP vaccine in mice, higher doses of RNAx potently suppress biomarkers of systemic reactogenicity while preserving the cellular and antibody response. At lower doses of RNAx, the antibody response was consistently enhanced after a single injection and occasionally following the boost. These findings establish RNAx as a modular and tunable platform technology for reducing reactogenicity while maintaining immunogenicity or in some contexts improving the antibody response of saRNA vaccines. The modular nature of encoding RNAx from a discrete mRNA unlocks the potential of improving any RNA-based vaccine, with the ability to fine tune dose for better tolerability and/or dose-sparing enabled by increases in potency.

## RESULTS

### RNAx enhances GOI expression from saRNA

RNAx, derived from the Cardiovirus leader protein, suppresses antiviral innate signaling^23^ and PKR-mediated inhibition of translation^24^, thereby enhancing gene expression ^29,30^ (Fig. 1A). We first combined RNAx with saRNA in two different forms: either as a single RNA molecule under the control of an internal ribosome entry site (IRES), “*in cis*”, or as a discrete mRNA, “*in trans*” (Fig. 1B). Using a strain of the Venezuelan equine encephalitis virus (VEEV) Simplicon vector, we cloned three reporter constructs under the control of the subgenomic promoter with RNAx *in cis* to estimate the impact on GOI expression: Influenza hemagglutinin (HA) fused at the C-terminus with nano-Luciferase (HA-nLuc), a secreted form of nLuc (sec-nLuc), and firefly luciferase (fLuc). We transfected these reporter constructs with or without RNAx *in cis* into BJ cells, a normal human diploid fibroblast cell line previously utilized for studying the impact of innate signaling on GOI expression from RNA^32^. Relative to saRNA only, saRNA with RNAx *in cis* enhanced the expression of HA-nLuc 62-fold, sec-nLuc 34-fold, and fLuc 8-fold (Fig. 1C). As a control for potential changes to the ability of IRES-containing constructs to express GOI, we also transfected 293T cells, which lack robust expression of innate sensors of dsRNA^33,34^. Each of the constructs expressed similar levels of reporter protein regardless of the presences of RNAx in 293T cells (Fig. 1D). When RNAx was co-transfected *in trans*, sec-nLuc expression was enhanced up to 11-fold in a dose dependent manner (Fig. 1E). We also transfected BJ cells with a nucleoside modified fLuc-encoding mRNA and treated cells with recombinant interferon-beta (IFN-β) or the dsRNA analog Poly(I:C), each of which suppressed fLuc expression over 10-fold relative to untreated cells (Fig. 1F). When co-transfecting cells with RNAx, fLuc activity increased by 4 to 7-fold in the presence of IFN-β or Poly(I:C), demonstrating that RNAx partially rescues translation in the presence of excessive innate signaling (Fig. 1B). Together, these results show that RNAx increases GOI expression from saRNA and rescues translation in the presence of excessive innate signaling.

**Figure 1.**
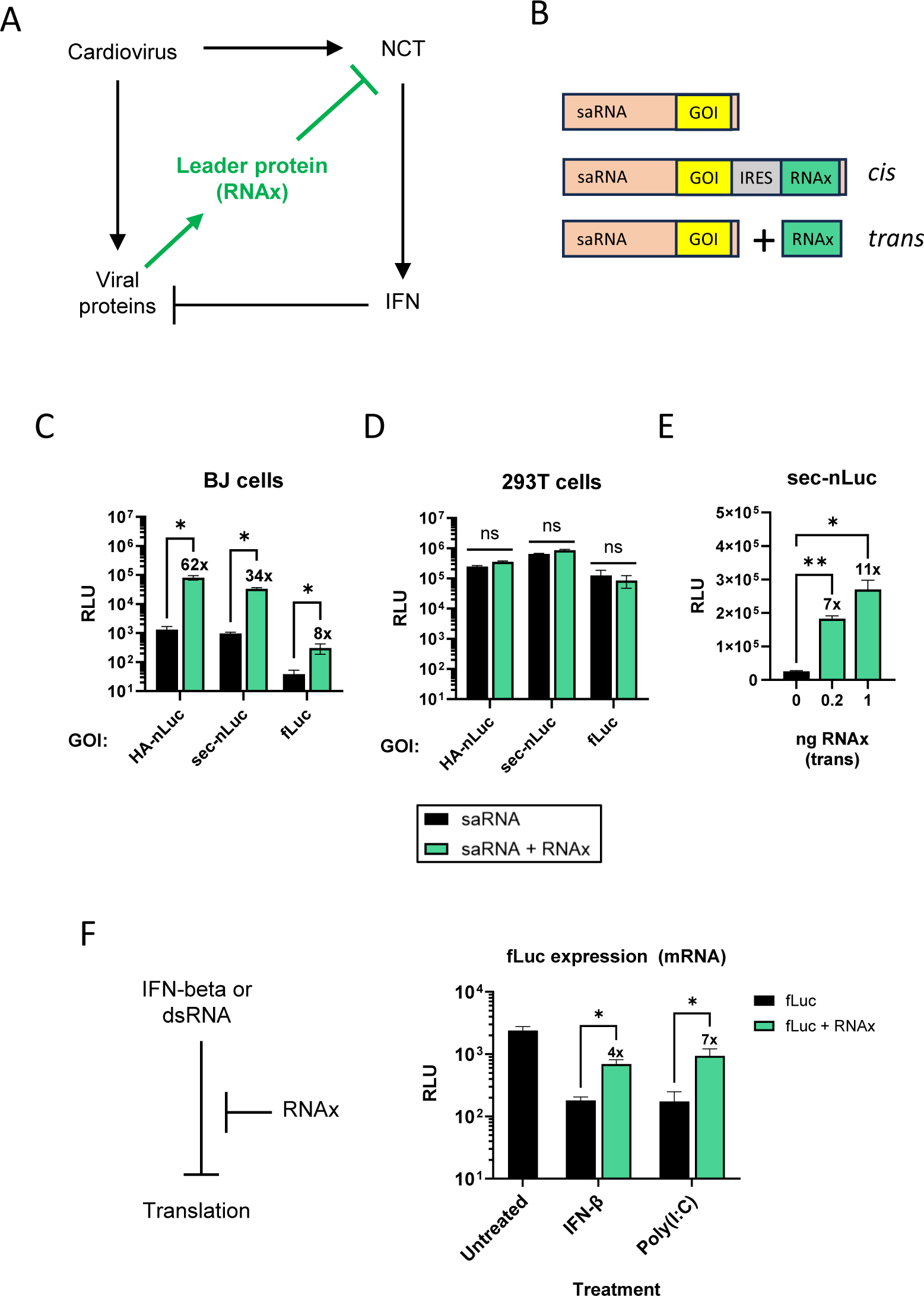
RNAx enhances GOI expression from saRNA *in vitro*. A) The Cardiovirus leader protein suppresses innate signaling by dampening nucleocytoplasmic transport (NCT) in infected cells. RNAx refers to an mRNA that encodes for the leader protein. B) RNAx was expressed from VEEV saRNA in cis from an IRES downstream of GOI or added in trans as an mRNA. C) BJ or D) 293T cells were transfected with reporter constructs and Luc signal measured 48hr later. E) RNAx was expressed in trans from a discrete mRNA alongside an saRNA expressing secreted nLuc saRNA. BJ cells were transfected with nLuc saRNA with increasing amounts of RNAx and 6 days post transfection supernatants were harvested for nLuc activity. F) RNAx reverses the inhibition of translation from RNA caused by IFN-β or dsRNA. BJ cells were transfected with a modified nucleoside fLuc-encoding mRNA and treated with either 1 ng/mL recombinant IFN-β or 100 ng/mL Poly(I:C). Luc activity was measured 48 hr post transfection. Mean ± SEM, n=3-4. *p<0.05, **p<0.01 Ratio paired t-test. Numbers above bars indicate average fold change enhancement of RNAx containing groups relative to saRNA only.

To determine whether RNAx enhances GOI expression from saRNA *in vivo*, we injected C57BL/6 mice intramuscularly with 2 µg of HA-nLuc saRNA-LNPs with or without RNAx *in cis* or *in trans* and monitored nLuc activity in vivo imaging system at days 1, 6, and 13 post injection (Fig. 2A-C). RNAx *in trans* yielded a statistically significant enhancement of 170-fold more nLuc expression relative to the saRNA alone group one day post injection (p<0.001, Kruskal-Wallis test with Dunn’s multiple comparisons) while changes induced by RNAx *in cis* were not statistically significant (Fig. 2D). Due to the high magnitude of GOI expression enhancement, modular nature, and ability to fine tune RNAx dose by co-formulating a distinct mRNA, we focused on RNAx *in trans* for future studies. Interestingly, nLuc expression in the groups receiving RNAx peaked at day 1 post injection while nLuc activity in the saRNA alone group peaked at day 6 (Fig. 2D). This is consistent with observations by others that GOI expression from saRNA^14^ or replication-incompetent RNA viruses^35,36^ peak in GOI expression at early time points post injection in mouse backgrounds deficient in IFN signaling. To examine whether RNAx could enhance GOI expression from another inflammatory RNA molecule in mice, we also tested whether RNAx modulated GOI expression from unmodified mRNA, an inducer of innate signaling *in vivo*^37^. RNAx increased GOI expression up to 12-fold 24 hr post injection (Supplementary Fig. 2A-B). These results demonstrate that RNAx enhances GOI expression from multiple forms of inflammatory RNA *in vivo*.

**Figure 2.**
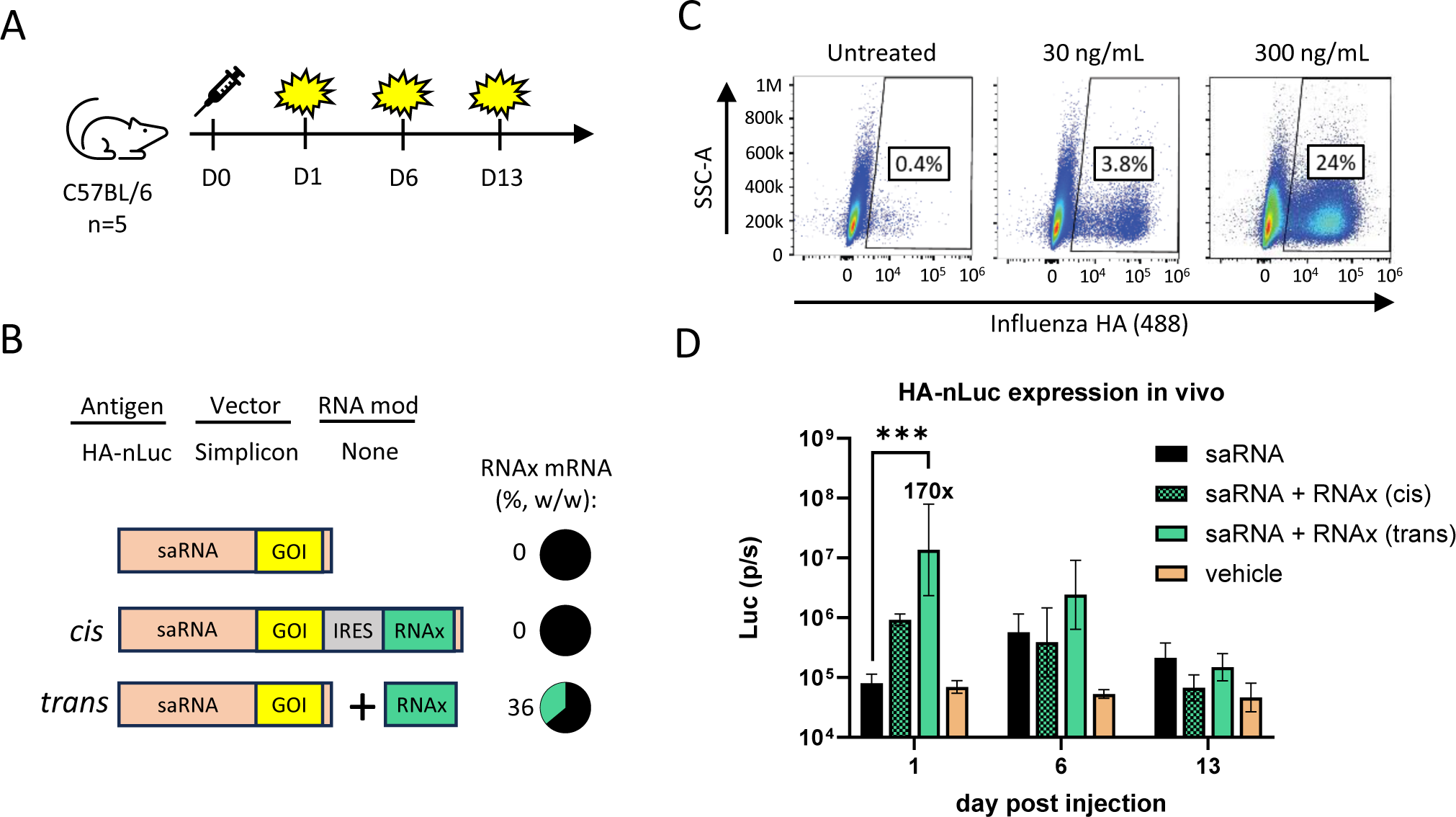
RNAx enhances GOI expression from saRNA *in vivo.* A) C57BL/6 mice (n=5 per group) were intramuscularly injected with 2 µg Influenza HA-nLuc expressing saRNA-LNPs and Luc activity at the injection site quantified with IVIS on the indicated days post injection. Total flux of Luc signal is represented by photons per sec (p/s). B) RNAx was co-delivered with Influenza HA-nLuc expressing saRNA either *in cis* or *trans* from a separate mRNA, the latter of which was co-formulated with 36% (weight/weight) RNAx. C) Flow cytometry of Influenza HA surface expression in 293T cells treated for 24hr with HA-nLuc saRNA-LNPs (no RNAx) at the indicated concentrations. D) Quantification of nLuc activity at the injection site quantified with IVIS on the indicated days post injection. Geometric mean ± Geometric SD, ***p<0.001, Kruskal-Wallis test with Dunn’s multiple comparisons. Numbers above bars indicate average fold change enhancement of RNAx containing groups relative to saRNA only.

### RNAx suppresses proinflammatory cytokines in primary human immune cells

Within its natural viral context, the Cardiovirus leader protein suppresses innate signaling pathways and IFN production^22,23^. We next quantified the impact of RNAx on the induction of proinflammatory cytokines and corresponding GOI expression from saRNA in primary human cells. We treated human peripheral blood mononuclear cells (PBMCs) with HA-nLuc saRNA-LNPs (Fig. 2C) and quantified the levels of 48 cytokines in the supernatants with a multiplex assay alongside GOI expression from lysates (Fig. 3A). HA-nLuc saRNA-LNPs without RNAx induced a total of 15 cytokines by at least 2-fold relative to untreated cells (p<0.05, ratio paired t-test), including IFN-α, IFN-γ, TNF-α, and IP-10 (Fig. 3B). With the inclusion of 36% RNAx mRNA with HA-nLuc saRNA cargo into LNPs, 14/15 of the saRNA-LNP-induced cytokines were suppressed at least 2-fold (p<0.05, ratio paired t-test) (Fig 3B). The most strongly suppressed cytokines were IFN-α (454-fold), MIP-1α (235-fold), and MCP-3 (137-fold) (Fig. 3B). We verified the levels of IFN-α secretion via ELISA and observed potent suppression with RNAx (Fig. 3C). We also observed a 2- to 6-fold enhancement of nLuc expression by RNAx compared to saRNA alone (Fig. 3D). To verify the specificity of RNAx-dependent suppression of IFN production based on its known mechanism of action, a zinc-finger domain mutant of RNAx, reported to completely inactivate the leader protein^22^, was unable to suppress the production of IFN-β in BJ cells treated with HA-nLuc saRNA-LNPs (Supplementary Fig. 1).

**Figure 3.**
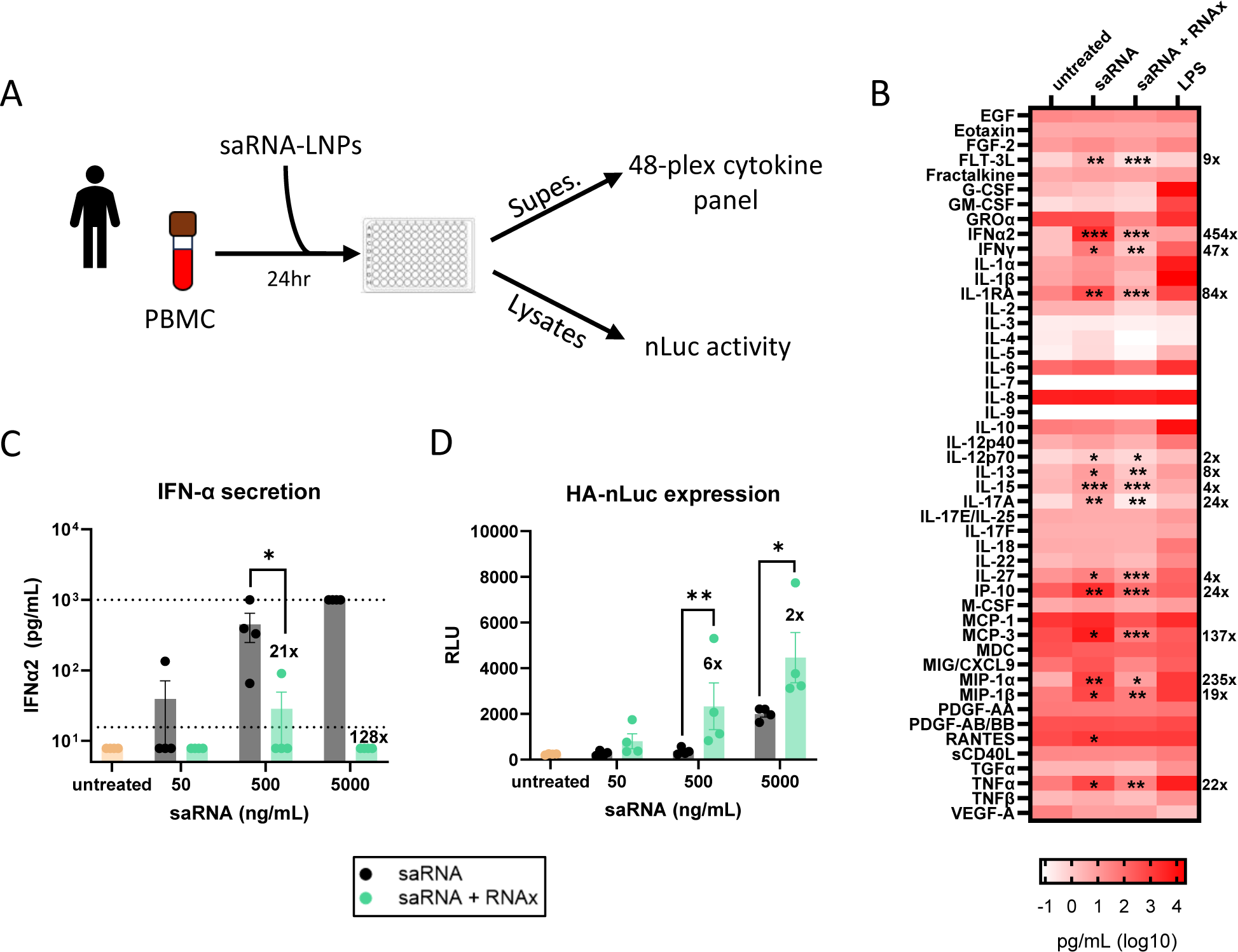
RNAx suppresses the expression of proinflammatory cytokines and enhances GOI expression in human PBMC. Human PBMC (n=4 donors) were treated with varying concentrations of HA-nLuc saRNA-LNPs for 24hr and supernatants harvested for cytokines and cell lysates for Luc activity. B) PBMCs treated with 5ug/mL saRNA-LNPs were analyzed for the abundance of a panel of human cytokines. Geometric mean. Asterisks in the ‘saRNA’ column indicate cytokines that were induced 2-fold on average by saRNA only vs. untreated cells (*p<0.05, **p<0.01, ***p<0.001 ratio paired t-test). Asterisks in the ‘saRNA + RNAx’ column indicate cytokines that were suppressed at least 2-fold on average by RNAx relative to saRNA only (*p<0.05, **p<0.01, ***p<0.001 ratio paired t-test), with the magnitude of the fold inhibition indicated to the right of the histogram. C) Supernatants from the same experiment were independently analyzed for IFN-α by ELISA and D) lysates for nLuc activity. Mean ± SEM, n=4 donors, *p<0.05, **p<0.01 Ratio paired t-test. Dotted lines = upper or lower limit of quantification. Numbers above bars indicate average fold change enhancement or inhibition of RNAx containing groups relative to saRNA only.

### RNAx suppresses serum biomarkers of reactogenicity from an Influenza HA saRNA-LNP vaccine

To test the hypothesis that RNAx suppresses proinflammatory cytokines linked to vaccine reactogenicity^38–40^ we designed an Influenza HA vaccine construct based on the TC-83 strain of VEEV by expressing only Influenza HA and employing 5-methylcytidine (5mC) modified nucleosides^17,18,41^. This construct robustly expressed HA in both 293T (Fig. 4A) and BJ cells (Supplementary Fig. 3A) and when co-formulated with RNAx (Fig. 4B), displayed enhanced HA expression in BJ cells (Supplementary Fig. 3B). Consistent with the suppression of primary IFN (IFN-α) in PBMCs (Fig. 3B-C), RNAx also suppressed the secretion of IFN-β and the expression of tetherin, a well-studied IFN-stimulated gene^42^ relative to HA saRNA alone in BJ cells (Supplementary Fig. 3C-E). Further supporting a link between innate signaling activation and GOI expression, saRNA did not induce IFN-β secretion or tetherin expression in 293T cells, alongside a lack of enhanced GOI expression from RNAx (Supplementary Fig. 3B-E).

**Figure 4.**
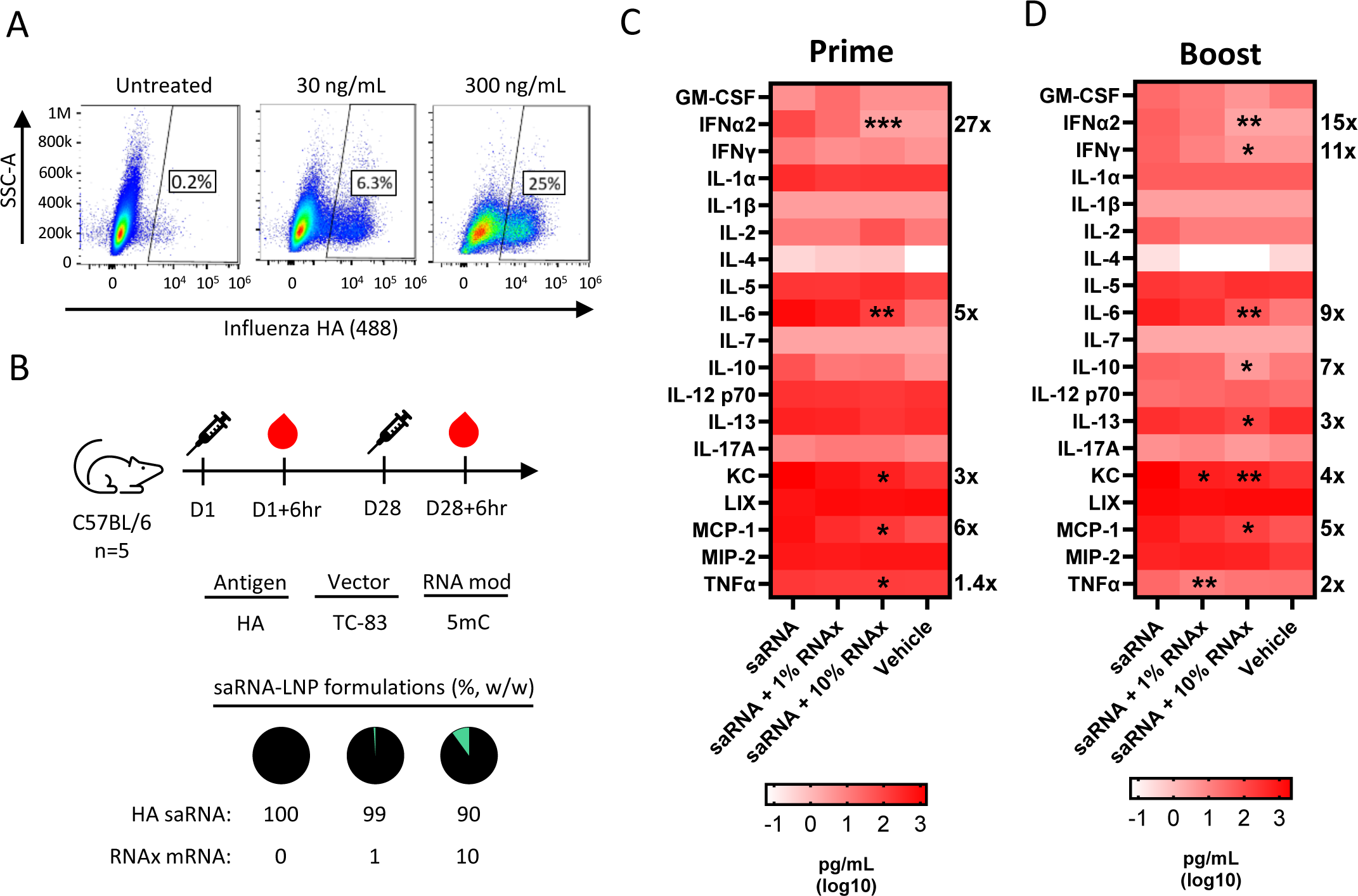
RNAx suppresses proinflammatory cytokines in mice following vaccination. A) C57BL/6 mice (n=5 per group) were intramuscularly injected with 0.5 µg of Influenza HA saRNA-LNPs and were analyzed for serum cytokines 6hr post injection following B) the prime or C) the boost at the indicated days of the study. Geometric mean. Asterisks indicate statistically significant reductions in cytokine levels (*p<0.05, **p<0.01, ***p<0.001, Kruskal-Wallis test with Dunn’s multiple comparisons) with the magnitude of the fold inhibition indicated to the right of each histogram. D) Flow cytometry of Influenza HA surface expression in 293T cells treated for 24hr with HA saRNA-LNPs (no RNAx) at the indicated concentrations.

C57BL/6 mice were vaccinated with 0.5 µg of 5mC-modified HA saRNA-LNP constructs at days 1 and 28 and serum was collected 6 hr after each injection (Fig. 4B). We quantified the levels of 18 proinflammatory cytokines with a multiplex assay and IFN-α with an ELISA. Changes in cytokine levels between groups were deemed significant at p<0.05 from a Kruskal-Wallis test with Dunn’s multiple comparisons. A summary of reactogenicity biomarker findings is presented in Table 1. Following the prime injection, RNAx suppressed the levels of five cytokines: IFN-α (27-fold), IL-6 (5-fold), KC (3-fold), MCP-1 (6-fold), and TNF-α (1.4-fold) (Fig. 4C, Table 1). Following the boost, RNAx suppressed levels of three other cytokines in addition to the five regulated by RNAx post prime: IFN-γ (11-fold), IL-10 (7-fold), and IL-13 (3-fold) (Fig. 4D, Table 1). Similar results were found from a vaccination study with the same HA saRNA-LNPs administered to BALB/c mice (Table 1, Supplementary Fig. 4) as well as an HA saRNA-LNP formulation that utilized the Simplicon vector that was used in earlier *in vitro* studies (Table 1, Supplementary Fig. 5). In the context of unmodified mRNA, RNAx suppressed levels of IFN-α (26-fold), IL-6 (5-fold), KC (3-fold), and MCP-1 (5-fold) (Supplementary Fig. 2C, Table 1). Of note, in nearly all cases, only groups with >8% RNAx (w/w) showed statistically significant inhibition of cytokine levels (Fig. 4C-D, Supplementary Fig. 4B-C, 5C-D). Across all saRNA studies IFN-γ, IL-6, MCP-1, and TNF-α were suppressed following the prime or boost in 2 out of 3 studies (Table 1), pointing to a robust and consistent suppression of cytokines by RNAx that correlate with vaccine adverse events in humans^38–40^.

**Table 1.** Summary of proinflammatory cytokine measurements. Compilation of results from Fig. 4 and Supplementary Fig. 4 and 4. Maximum fold change suppression from RNAx containing saRNA-LNPs relative to saRNA only LNPs with p<0.05 (Kruskal-Wallis test with Dunn’s multiple comparisons). Empty cells reflect differences that were not statistically significant. n.t. = not tested.

|  | Prime |  |  |  | Boost |  |  |
| --- | --- | --- | --- | --- | --- | --- | --- |
| Vector | TC-83 | TC-83 | Simplicon | mRNA | TC-83 | TC-83 | Simplicon |
| RNA mod | 5mC | 5mC | Natural | Natural | 5mC | 5mC | Natural |
| Mouse strain | C57BL/6 | BALB/c | C57BL/6 | C57BL/6 | C57BL/6 | BALB/c | C57BL/6 |
| GM-CSF |  |  |  |  |  |  |  |
| IFN-α2 | 27x | 20x | 7x | 26x | 15x | 28x | 11x |
| IFN-γ | | 11x | 41x | | 11x | 15x | |
| IL-1α | | | n.t. | | | | |
| IL-1β | | | n.t. | | | | |
| IL-2 |  | 3x |  |  |  |  |  |
| IL-4 |  |  |  |  |  |  |  |
| IL-5 |  |  | n.t. |  |  |  |  |
| IL-6 | 5x | 9x | 14x | 5x | 9x |  | 4x |
| IL-7 |  |  | n.t. |  |  |  |  |
| IL-10 |  |  |  |  | 7x |  |  |
| IL-12 p70 |  |  |  |  |  |  |  |
| IL-13 |  |  | n.t. |  | 3x |  |  |
| IL-17A |  |  | n.t. |  |  |  |  |
| KC | 3x |  | n.t. | 3x | 4x |  | 3x |
| LIX |  |  | n.t. |  |  |  |  |
| MCP-1 | 6x | 7x | 5x | 5x | 5x | 6x |  |
| MIP-2 |  |  | n.t. |  |  |  |  |
| TNF-α | 1.4x | 3x | | | 2x | | 1.5x |

### RNAx does not alter the magnitude or Th1/Th2 balance of the cellular response

Because cellular response to vaccination is heavily influenced by the accompanying induction of innate signaling^7^, we next tested whether the potent suppression of the innate response by RNAx in turn negatively impacted the cellular response to vaccination. C57BL/6 mice were vaccinated at day 1 and 28 with 0.5 µg or 0.05 µg of HA saRNA-LNPs (TC-83 vector) with increasing amounts of co-formulated RNAx mRNA (Fig. 5A). Two weeks post boost, splenocytes were isolated and stimulated with an overlapping pool of HA-derived peptides for quantification of IFN-γ and IL-4 secreting cells by ELISPOT. Among the saRNA alone groups receiving 0.5 µg or 0.05 µg doses, only the 0.5 µg dose group yielded a statistically significant increase in IFN-γ-secreting cells vs. vehicle only (p=0.0011, Kruskal-Wallis test with Dunn’s multiple comparisons), whereas IL-4 responses remained unchanged compared to vehicle at both doses. Within all dose groups, the inclusion of RNAx did not alter the number of IFN-γ (Fig. 5B) or IL-4 secreting cells (Fig. 5C). Identical results were obtained when examining splenocytes from mice vaccinated with 0.5 µg of the Simplicon-derived HA saRNA-LNP formulations (Supplementary Fig. 7A-C). Together, these data show that the overall balance as well as the Th1-biased nature of the cellular response is preserved with the inclusion of RNAx.

**Figure 5.**
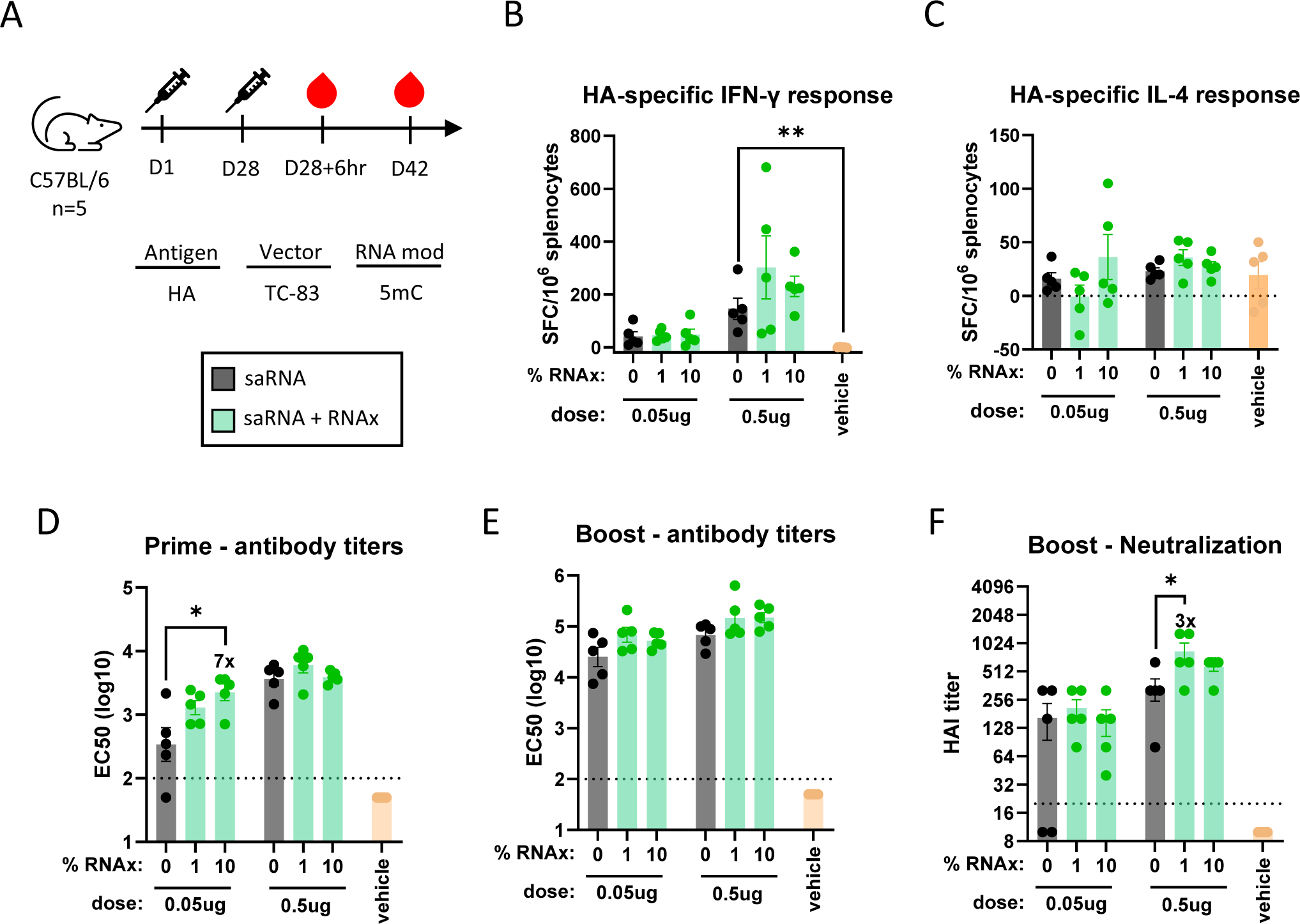
Impact of RNAx on immunogenicity from an Influenza HA saRNA-LNP vaccine. A) C57BL/6 mice (n=5 per group) were intramuscularly injected (prime and boost) with 0.05 µg or 0.5 µg of HA saRNA-LNPs. Serum was collected 6hr after the boost (post prime measurement) and serum and splenocytes collected 2 weeks post boost. Splenocytes were treated with 1 ug/mL Influenza HA peptide pools and analyzed for B) IFN-γ and C) IL-4 secreting cells by ELISPOT. Spot forming cells (SFC) were subtracted by untreated samples and normalized by 10^6^ splenocytes. Mean ± SEM. Total anti-HA IgG in the serum (EC50) was quantified by ELISA at D) post prime and E) 2 weeks post boost. F) Neutralization titers were quantified by HAI assay 2 weeks post boost. Dotted line = lower limit of quantification, Geometric mean ± Geometric SD, *p<0.05, Kruskal-Wallis test with Dunn’s multiple comparisons. Numbers above bars indicate fold enhancement of RNAx containing groups relative to saRNA only.

### RNAx enhances antibody titers after single injection of HA saRNA-LNP vaccine

The induction of IL-6 has been shown to be a potent adjuvant for the antibody response to RNA-LNP vaccines^43^. Given that RNAx potently suppressed IL-6 and other cytokine levels post prime and boost (Table 1), we next measured the impact of RNAx on the antibody response. C57BL/6 mice were vaccinated at day 1 and 28 with 0.5 µg or 0.05 µg of 5mC-modified TC-83 HA saRNA-LNPs with 1% or 10% co-formulated RNAx (Fig. 5A). Four weeks after the prime, serum was analyzed for anti-HA binding IgG antibodies by ELISA. At the 0.05 µg dose, we observed a statistically significant 7-fold increase in binding antibody titers with 10% RNAx (p<0.05, Kruskal-Wallis test with Dunn’s multiple comparisons) (Fig. 5D), while at a 0.5 µg total dose, antibody levels remained constant regardless of RNAx (Fig. 5D). In vaccinated BALB/c mice, we observed a 3-fold increase (p<0.05, Kruskal-Wallis test with Dunn’s multiple comparisons) in binding antibody titers post prime from 1% RNAx at the 0.05 µg total dose (Supplementary Fig. 6A-B). We also examined the antibody response in C57BL/6 mice vaccinated with the Simplicon vector HA saRNA-LNP constructs (Supplementary Fig. 5A-B, 6A) and found that the 2% RNAx group showed a statistically significant (p<0.05, Kruskal-Wallis test with Dunn’s multiple comparisons) 7-fold enhancement, while titers from groups with higher amounts of RNAx returned to levels of the control saRNA group (Supplementary Fig. 7D). While the optimal % of RNAx in the formulation for enhancing antibody titers varied across mouse strains and VEEV vectors, in no condition tested did RNAx suppress the antibody response.

### RNAx preserves and occasionally enhances the antibody response after vaccine boost

C57BL/6 mice vaccinated with 5mC-modified TC-83 HA saRNA-LNPs were also analyzed for the presence of binding antibody and viral neutralizing titers (HAI assay) 2 weeks post boost. At the 0.5 µg total dose, neutralization titers were enhanced 3-fold (p<0.05, Kruskal-Wallis test with Dunn’s multiple comparisons) with 1% RNAx (Fig. 5F). While binding antibody titers increased 2-fold with this group, this difference was not statistically significant (Fig. 5E).

However, in BALB/c mice vaccinated with a 0.05 µg dose, a dose with equivalent binding titers as 0.5 µg vaccinated C57BL/6 mice, 1% RNAx did enhance anti-HA antibody titers 2-fold in a statistically significant manner (p<0.05, Kruskal-Wallis test with Dunn’s multiple comparisons) (Supplementary Fig. 6C). Neutralization titers in BALB/c mice receiving 1% RNAx at the 0.05 µg total dose increased slightly but failed to reach statistical significance (p=0.067, Kruskal-Wallis test with Dunn’s multiple comparisons) (Supplementary Fig. 6C). As with the antibody response following the prime, conditions that yielded enhancements to the antibody response varied across mouse strains and VEEV vectors. Crucially however, in all other conditions tested, including groups receiving 10% RNAx, binding antibody and neutralization titers were preserved (Fig. 5E-F, Supplementary Fig. 6C-D, 6E-F), despite potent suppression of serum biomarkers of innate signaling (Table 1).

In conclusion, our results cumulatively show that by incorporating RNAx into saRNA vaccines of various backbones and nucleotide modifications, serum biomarkers of vaccine reactogenicity are greatly reduced, in many cases close to the baseline. Despite suppressing innate signaling and reactogenicity parameters that correlate with traditional avenues of adjuvancy, RNAx did not negatively affect immunogenicity. In fact, in certain conditions, RNAx enhanced the antibody response. This suggests that RNAx can be used to control RNA vaccine side effects while preserving the immune response or enable dose sparing via increased per µg potency.

## DISCUSSION

Reducing saRNA-induced innate signaling is an active area of research for improving patient tolerability. One approach has been to incorporate modified nucleotides into the saRNA molecule. Relative to unmodified nucleosides, modified nucleoside-containing saRNA vaccines have been shown to reduced serum levels of IFNα 2.4-fold in mice^17^ and may be associated with lower incidence of systemic adverse events in the clinic^41^. RNAx builds upon the results by further suppressing IFNα levels to below the limit of quantification in our assays, as well as potently and consistently reducing levels of IFN-γ, IL-6, MCP-1, and TNF-α. We hypothesize that RNAx-dependent suppression of these cytokines will improve saRNA vaccine tolerability. In humans, serum levels of IFN-γ, IL-6 and MCP-1 correlated with systemic symptoms following boost with BNT162b2^38^ while in the context of an adjuvanted hepatitis B virus vaccine, IFN-γ and IL-6 correlated with systemic symptoms following boost^39^. In addition, serum levels of TNF-α correlated with injection site soreness following injection with an attenuated Influenza vaccine^40^. Supporting these clinical data, IFN-α^44^, IL-6^45^, and TNF-α^46^ are well known to induce flu-like symptoms when administered systemically in humans. While there is a paucity of clinical data quantifying the association of serum biomarkers with saRNA-LNP vaccine tolerability and immunogenicity, future clinical studies deploying RNAx could directly address this question.

Optimal immunogenicity is a fine balance between innate signaling (adjuvancy) and antigen expression^7^. Intuitively, one would expect that suppressing innate signaling-driven reactogenicity would also reduce immunogenicity. In the case of RNAx, despite potent suppression of biomarkers of innate signaling in mice and human PBMC, RNAx did not suppress immunogenicity at any condition tested. We propose that this is due to suppression and not elimination of all necessary innate signaling in conjunction with a corresponding increase in antigen expression. In addition, RNAx suppressed IFN-α and MCP-1, molecules linked to pathways that can suppress immunogenicity^14,47,48^. At lower doses of RNAx that did not potently suppress serum proinflammatory cytokines, we tended to detect enhancements to the antibody response. This highlights a versatile feature of RNAx in that different doses of RNAx may be optimal for different contexts. For example, one could either improve the tolerability of an already highly potent saRNA-LNP vaccine or increase the per µg potency, thereby facilitating dose sparing.

While we obtained evidence of RNAx enhancing the antibody response in some contexts, mice are well-known to underestimate the positive effects on vaccine immunogenicity when modulating innate signaling from RNA-LNPs. Nucleoside modifications^34,37^ were instrumental in enabling the first clinically approved SARS-CoV-2 mRNA-LNP vaccines BNT162b2^49^ and mRNA-1273^50^; however, an unmodified mRNA-LNP vaccine developed by CureVac (CVnCoV) that failed in the clinic^51^ generated similar levels of immunogenicity in mice compared to the modified nucleoside vaccines^52–54^. Bernard and colleagues compared modified and unmodified nucleoside versions of an mRNA-LNP vaccine directly in mice and non-human primates (NHP) and found that mice greatly underestimated the beneficial effects of modified nucleosides^55^. CureVac’s next generation unmodified mRNA platform (CV2CoV) was engineered to have more efficient antigen expression, which not only outperformed CVnCoV but also achieved similar immunogenicity with BNT162b2 in NHPs^56^. However, CVnCoV and CV2CoV yielded comparable immunogenicity results in mice^57^. Mirroring our findings that RNAx occasionally enhanced immunogenicity in mice, perturbation of the IFN pathway has been found to either enhance^14^ or not impact^11,12,20^ the antibody response from saRNA-LNP vaccines in mice. In total, the body of work from RNA-LNP vaccines suggest caution is warranted when interpreting vaccine improvements that dampen innate signaling based solely on mouse immunogenicity studies. We propose a holistic approach that incorporates mechanistic studies both in primary human cells as well as mice. Future studies in larger animals or humans are ideally suited to uncover the true impact of RNAx on vaccine immunogenicity.

Co-delivering RNAx *in trans* is an innovative approach to balancing saRNA-driven innate signaling by virtue of being broadly acting, modular, and tunable. There is growing evidence that saRNA platforms activate excessive innate signaling^6^ and taming this activation is an active area of research^11,12,15–18,20,21^. One approach has been to co-express inhibitors of innate signaling pathways *in cis*. Co-delivering the MERS ORF4a protein *in cis*, a suppressor of IFN production and PKR activation^58^, enhanced antibody titers in rabbits but not mice from a Rabies virus vaccine^11^. Inclusion of the Vaccinia virus proteins ‘EKB’, inhibitors of the IFN pathway and PKR activation, *in cis* was shown to enhance GOI expression^16^, but any effect on immunogenicity has not been made public to our knowledge. RNAx is distinct from these approaches in that it acts broadly by targeting NCT, not individual innate signaling pathways, and because of its modular nature, which enables application of different doses for any saRNA without having to modify the antigen expressing saRNA molecule. We also focused our efforts on evaluating the role of RNAx only in the context of LNP-based delivery as a proof of principle. A study by Kimura and colleagues showed that transient perturbation of the IFN pathway through co-delivery of a primary IFN receptor (IFNAR) blocking antibody reliably enhanced antibody titers of an saRNA vaccine with their lipid inorganic nanoparticle (LION) platform, but not with LNPs^12^. Given its potent suppression of IFN, we hypothesize that RNAx will also enhance the antibody response from next generation delivery platforms such as LION and nanostructured lipid carrier (NLC)^12,59^.

In conclusion, our data demonstrate that co-delivery of RNAx is an effective means of dampening excessive innate signaling driven by saRNA vaccines without sacrificing immunogenicity. Supported by our data with saRNA and unmodified mRNA, any reactogenic nucleic acid-based therapy should benefit from RNAx. For vaccines, we hypothesize that suppressing saRNA-driven innate signaling will improve tolerability as well as enhance immunogenicity for any saRNA platform or target in clinic.

## METHODS

### Cells

Cells were passaged at 37°C in 5% CO_2_. BJ fibroblasts (ATCC, CRL-2522) and 293T/17 cells (ATCC, CRL-11268) were passaged in complete DMEM (Gibco 10566-016), containing 10% heat-inactivated FBS (VWR, 89510-186) and 100U/mL of Penicillin-Streptomycin (Gibco, 72400-047).

### Cloning and *in vitro* transcription of RNA

saRNA constructs encoding nLuc, fLuc, the Influenza HA-nLuc fusion, and Influenza HA were derived from the Simplicon vector (Sigma, SCR724). Venezuelan equine encephalitis virus (VEEV) sequences were amplified by PCR and cloned using HiFi assembly (NEB, E2621).

Secreted nLuc was engineered by fusing the human albumin signal peptide (MKWVTFISLLFLFSSAYS) to the N-terminus of nLuc. Influenza HA amino acid sequence was derived from accession ACP41953.1. The TC-83 strain of VEEV (Accession MZ399798.1) was also cloned to express Influenza HA (ACP41953.1).

In vitro transcription (IVT) of nLuc, fLuc, or Influenza HA-nLuc saRNA constructs was conducted in-house. IVT templates were generated using PCR with primers encoding a 30 nt polyA tail. The PCR products were gel purified (Zymo research, D4008), followed by an additional DNA purification (NEB, T1030). IVT reactions (NEB, E2040S) were carried out with natural nucleosides and co-transcriptional capping with CleanCap AU (Trilink, N-7114). RNA was purified with silica membrane spin columns (NEB, T2050) and quantified by NanoDrop. In vitro transcription of the Influenza HA saRNA construct derived from the Simplicon vector was performed by Aldevron (Fargo, USA) with natural nucleosides and co-transcriptional capping with CleanCap AU (Trilink, N-7114). In vitro transcription of the Influenza HA saRNA construct derived from TC-83 was performed by Trilink (San Diego, USA) using 5-Methylcytosine (5mC) and co-transcriptional capping with CleanCap AU (Trilink, N-7114). N1-Methylpseudouridine-5’-Triphosphate-modified, CleanCap-AG RNAx mRNA as well as the size-matched, noncoding filler mRNA were synthesized by Trilink (San Diego, USA). Unmodified (L-7602) and 5moU-modified (L-7202) fLuc-encoding mRNA was obtained from Trilink (San Diego, USA).

### Lipid nanoparticle (LNP) formulations

LNPs were prepared by microfluidic mixing as follows. First, the RNA samples were diluted to a final concentration of 50 µg/mL in 50 mM NaOAc pH 4.5 (Thermo Scientific Chemicals J63669AK) and 150 mM NaCl (Thermo Scientific Chemicals J60434AE). The lipid mix was prepared in 100% ethanol (Fisher BioReagents BP2818100) by diluting ALC-0315 ionizable lipid (Cayman Chemicals 34337), DSPC (Cayman Chemicals 15100), cholesterol (Cayman Chemicals 9003100), and ALC-0159 PEGylated lipid (Cayman Chemicals 34336). For saRNA-LNPs, lipids were used at the molar ratio of 46.3% ALC-0315 / 9.4% DSPC / 43.5% cholesterol / 0.8% ALC-0159 at a final total lipid concentration of 8.09 mM. For fLuc mRNA-LNPs, lipids were used at the molar ratio of 46.3% ALC-0315 / 9.4% DSPC / 42.7% cholesterol / 1.6% ALC-0159 at a final total lipid concentration of 8.09 mM. Before each mixing reaction, the flow channel of the microfluidic mixing chip was primed by performing a mock mixing reaction using 1X formulation buffer (without cargo) and 100% EtOH (without lipids) in a total volume of 2 mL. RNA and the lipid mix were combined using the Flex-M microfluidic mixing system (Precigenome) using an RNA-to-lipid flow rate ratio (FRR) of 3:1 at the total flow rate (TFR) of 3 mL/min. Immediately after the mixing, the formed LNPs were diluted 10X in 1X Tris-buffered saline pH 7.4 (Fisher BioReagents BP2471-100) and incubated at RT for 1 hour. Thereafter, the stabilized LNPs were concentrated to 0.5 mL and washed 2X using Tris-buffered saline pH 7.4 using the Amicon Ultra 10kDa MWCO centrifugal filter units (EMD Millipore UFC901024).

Sucrose (Thermo Scientific Chemicals AC419762500) in Tris-buffered saline pH 7.4 was added to the LNP formulations as a cryoprotectant at the final concentration of 10% before storage at −80°C. Sample concentration and encapsulation efficiency were determined using the Quant-iT Ribogreen Assay (Invitrogen R11491). LNP Z-average size and polydispersity index (PDI) were measured using dynamic light scattering (Zetasizer, Malvern). A summary of QC measurements for LNP formulations are listed in Supplementary Table 1.

### *In vitro* transfection

BJ cells were seeded into tissue culture treated-white or standard 96-well cell culture plates in complete DMEM and incubated overnight at 37°C and 5% CO_2_. Opti-MEM cell culture media (Gibco, 31985-062) and Mirus TransIT mRNA-transfection reagent (Mirus Bio, MIR 2225) was brought to room temperature. All mixes were prepared under the biosafety cabinet and all surfaces and pipets were cleaned with RNAse Zap prior to handling of RNA. RNA was thawed on ice and kept on ice during the entire procedure. Up to 100 ng per well of RNA was mixed with Mirus transfection reagent according to the manufacturer’s instructions. If applicable, transfection complexes were added to cells immediately before adding 100 ng/mL Poly(I:C) (Fisher Scientific, 42-871-0) or 1 ng/mL IFN-β (Fisher Scientific, 84-99I-F010) and incubated at 37°C and 5% CO_2_. nLuc (Promega, N1120) or fLuc (Promega, G7940) activity was measured two to six days post transfection from the lysates (fLuc, HA-nLuc) or supernatants (secreted nLuc) according to the manufacturer’s instructions. Plates were then read on the Biotek Synergy LX plate reader for luminescence. IFN-β protein levels were quantified from supernatants by ELISA (R&D Systems, DIFNB0) or Lumit assay (Promega, CS2032C03). Values below the lower limit of quantification (LOQ) were transformed to 0.5*LOQ.

### Human PBMC

Cryopreserved human PBMCs were obtained from IQ Biosciences (cat# IQB-PBMC102). Donor characteristics are listed in Supplementary Table 2. Cells were thawed at 37°C, pelleted, and resuspended in prewarmed RPMI 1640 (Gibco 61870-036) containing heat-inactivated FBS (VWR, 89510-186). 50 µL of 5×10^6^ cells/mL were added to V-bottom 96-well plates and mixed with 50 µL of HA-nLuc saRNA-LNPs for a final concentration of 5,000 ng/mL, 500 ng/mL, or 50 ng/mL. Untreated and LNP-treated cells (100 pg/mL final, Invitrogen, 00-4976-93) served as controls. 24 hr later, cells were spun at 300g for 5 mins and supernatants stored at −80°C until cytokine measurements. Cell pellets were analyzed for nLuc activity (Promega, N1120) according to manufacturer’s instructions. Lysates were transferred to opaque white plates and read on the Biotek Synergy LX plate reader.

Undiluted cell supernatants were analyzed for the presence of 48 human cytokines using the Luminex™ 200 system (Luminex, Austin, TX, USA) by Eve Technologies Corp. (Calgary, Alberta). Forty-eight markers were simultaneously measured in the samples using Eve Technologies’ Human Cytokine Panel A 48-Plex Discovery Assay (Millipore Sigma, Burlington, Massachusetts, USA) according to the manufacturer’s protocol. The 48-plex consisted of sCD40L, EGF, Eotaxin, FGF-2, FLT-3 Ligand, Fractalkine, G-CSF, GM-CSF, GROα, IFN-α2, IFN-γ, IL-1α, IL-1β, IL-1RA, IL-2, IL-3, IL-4, IL-5, IL-6, IL-7, IL-8, IL-9, IL-10, IL-12(p40), IL-12(p70), IL-13, IL-15, IL-17A, IL-17E/IL-25, IL-17F, IL-18, IL-22, IL-27, IP-10, MCP-1, MCP-3, M-CSF, MDC, MIG/CXCL9, MIP-1α, MIP-1β, PDGF-AA, PDGF-AB/BB, RANTES, TGFα, TNF-α, TNF-β, and VEGF-A. Cytokines were deemed to be induced by saRNA-LNP if the geometric mean levels were 2-fold higher than untreated and p<0.05 by the ratio paired t-test. Cytokines were deemed suppressed by RNAx if the geometric mean levels were 2-fold lower in the RNAx groups relative to saRNA only groups and p<0.05 by the ratio paired t-test. IFN-α2 levels were independently quantified from the same cell supernatants with the LumiKine Xpress hIFNa 2.0 ELISA (InvivoGen luex-hifnav2) according to the manufacturer’s instructions. Values below the lower limit of quantification (LOQ) were transformed to 0.5*LOQ.

### Flow cytometry

3×10^5^ 293T cells/well and 1×10^5^ BJ cells/well were seeded into 12-well plates in complete DMEM overnight at 37°C and 5% CO_2_. LNPs were diluted in prewarmed complete DMEM and incubated with cells for 24 hr at 37°C and 5% CO_2_. Supernatants were collected and frozen at −80°C while cells were treated with trypsin (Gibco, 12605-010) for 1-5 mins, rinsed in complete DMEM, washed in PBS(-/-) (Gibco, 14190-136), then transferred to V-bottom 96-well plates. Cells were then incubated with LIVE/DEAD cell dye for 10 min at RT in the dark. Cells were rinsed in FACS buffer (PBS(-/-) + 1% heat inactivated FBS + 0.05% sodium azide) and then incubated with primary antibody for 1 hr at 4°C in the dark. Human anti-Influenza antibody clone FI6V3 (Creative Biolabs, PABL-214) was conjugated with the Zenon human IgG labeling kit according to manufacturer’s instructions (Invitrogen, Z25402). Anti-tetherin (CD317)-APC clone RS38E antibody (Biolegend, 348410) was used to stain for tetherin expression. Cells were rinsed twice in FACS buffer then fixed in 1% PFA (Electron Microscopy Sciences, 15714-5) for 1 hr at 4°C. For compensation beads, UltraComp eBeads Plus Compensation Beads (Fisher Sci, 01-333-342) were used for antibody complexes and ARC amine reactive kit (Fisher Sci, 50-112-1496) for LIVE/DEAD dye. Samples were run on the Attune NxT flow cytometer (Thermo Fisher) and analyzed with FlowJo (10.10.0).

### Animal studies

Animal studies at Charles River Laboratories (Worcester, MA) were conducted in compliance with CRL IACUC under IACUC No. I035. Animal studies performed at Labcorp (Denver, PA) were in compliance with the U.S. Department of Agriculture’s (USDA) Animal Welfare Act (9 CFR Parts 1, 2, and 3); the Guide for the Care and Use of Laboratory Animals (Institute of Laboratory Animal Resources, National Academy Press, Washington, D.C., 2011); and the National Institutes of Health, Office of Laboratory Animal Welfare. Whenever possible, procedures in these studies were designed to avoid or minimize discomfort, distress, and pain to animals. Studies conducted at Labcorp were conducted in accordance with Labcorp IACUC protocols, Standard Operating Procedures (SOP), Quality Management Systems (QMS), and Quality Principles (QP).

### In vivo imaging

*In vivo* imaging was performed at Charles River Laboratories (CRL, Worcester, MA) using 8-10 week old female C57BL/6 mice obtained from CRL. Mice were housed in Innovive individually ventilated cages kept under positive pressure during the study and acclimated for 2 days prior to injection. Test articles were thawed and stored at 4°C. Within 30 min of injection, test articles were equilibrated to room temperature. Mice were injected intramuscularly into the right quadricep with 50 µL containing 2 µg of saRNA-LNPs, 5 µg of mRNA-LNPs, or 50 µL of 10% sucrose in Tris-buffered saline pH 7.4 (vehicle). On days 1, 6, and 13 post injection, nLuc signal was quantified within 2 hr following 50 µL intraperitoneal injection of reconstituted Nano-Glo in Vivo FFz substrate (Promega, CS320501) as photons per sec (p/s) using the Lumina III *in vivo* imaging system (IVIS) (Perkin-Elmer). Data were analyzed in Prism 10.4.0 (GraphPad).

### Vaccination studies

Immunogenicity and reactogenicity studies were performed at Labcorp (Denver, PA). 10-12 week old female C57BL/6 or BALB/c mice obtained from CRL. Test articles were thawed and stored at 4°C. Within 30 min of injection, test articles were equilibrated to room temperature.

Mice were injected intramuscularly into the right quadricep with 50 µL containing 2 µg, 0.5 µg, or 0.05 µg of saRNA-LNPs or 50 µL of 10% sucrose in Tris-buffered saline pH 7.4 (vehicle) at the indicated days per study. Mice were bled retro-orbitally and processed into serum. Samples from animal studies examining immunogenicity read outs were also used for reactogenicity (serum cytokine) measurements.

### In vivo cytokine measurements

Serum was diluted 1:2 or 1:6 in PBS and analyzed for cytokine, chemokine and growth factor protein levels using the Luminex 200 system (Luminex, Austin, TX, USA) at Eve technologies Corp. (Calgary, Alberta). Samples from animals injected (primed) with 2 µg of the Simplicon saRNA vector were analyzed for the presence of ten markers using Eve Technologies’ Mouse Focused 10-Plex Discovery Assay (MilliporeSigma, Burlington, Massachusetts, USA) according to the manufacturer’s protocol. The 10-plex consisted of GM-CSF, IFNγ, IL-1β, IL-2, IL-4, IL-6, IL-10, IL-12p70, MCP-1, and TNFα. All remaining in vivo samples were analyzed for the presence of eighteen markers using Eve Technologies’ Mouse High Sensitivity 18-Plex Discovery Assay (MilliporeSigma, Burlington, Massachusetts, USA) according to the manufacturer’s protocol. The 18-plex consisted of GM-CSF, IFNγ, IL-1α, IL-1β, IL-2, IL-4, IL-5, IL-6, IL-7, IL-10, IL-12(p70), IL-13, IL-17A, KC/CXCL1, LIX, MCP-1, MIP-2 and TNFα. IFN-α2 levels were quantified with the LumiKine Xpress hIFNa 2.0 ELISA (InvivoGen luex-hifnav2) according to the manufacturer’s instructions. Values below the lower limit of quantification (LOQ) were transformed to 0.5*LOQ. Histograms show geometric means, with differences in abundance from RNAx deemed statistically significant if p<0.05 with the Kruskal-Wallis test with Dunn’s multiple comparisons.

### Anti-Influenza HA ELISA

96-well plates (Greiner BioOne, 655061) were coated with 1 µg/mL of recombinant Influenza HA antigen (Sino Biological, 11085-V08H-100) in PBS overnight at 4°C. Plates were washed 3x with wash buffer (PBS-T + 0.05% Tween-20) with a BioTek 405LS plate washer and incubated with EZ Block blocking buffer (ScyTek, EZB999) for 2 hr at 37°C. Plates were then washed 3x with wash buffer and incubated with diluted, heat-inactivated (56°C for 30min) mouse serum for 1 hr at RT. Samples were run in technical duplicate. Plates were then washed 3x with wash buffer and incubated with a 1:4,000 dilution (in EZ block) of Goat Anti-Mouse IgG Fc-HRP secondary (Southern Biotech, 1033-05) for 1 hr at RT. Plates were then washed 5x with wash buffer and incubated in pre-warmed (37°C) BioFx TMB One Component HRP Microwell Substrate (Surmodics, TMBW-1000-01) for 30 min at RT, then BioFx 450 nm Liquid Stop Solution for TMB Microwell Substrate (Surmodics, LSTP-100-01) added to plates and read on the Biotek Synergy LX plate reader at absorbance 450 nm within 5 mins. Technical replicate absorbance values were averaged and EC50 curves generated with the log(agonist) vs. response -- Variable slope (four parameters) regression analysis with a top=4 constraint in Prism (Version 10.4.0). Values below the lower limit of quantification (LOQ) were transformed to 0.5*LOQ. Data are presented as geometric mean ± geometric SD and statistical significance deemed if p<0.05 (Kruskal-Wallis test with Dunn’s multiple comparisons).

### ELISPOT

Spleens were removed aseptically and placed in a sterile 5 mL tube containing 3.5 mL of 4C CTL media (CTLT-010, CTL) and stored at 2-8°C until processing. Within 2hr, spleens and media were transferred to a gentleMACS C tube (130-093-237, Miltenyi Biotec) and placed into the gentleMACS Octo Dissociator (Miltenyi Biotec). Spleens were then homogenized using the program “m_spleen_04_01” and homogenates spun at 300g for 8 min. Supernatant was removed and red blood cells lysed with ACK lysis buffer (A10492-01, Gibco) for 4 min at RT. Lysis was neutralized with 2 mL of CTL media and cells spun at 300g for 8 mins. Supernatant was gently removed and cells resuspended in 5 mL CTL media + 1% Glutamax (35050-061, Gibco) + 1% penn/strep (Sigma, P4333). Cell suspension was passed through a sterile 70 µm tissue screen and counted.

IFN-γ and IL-4 producing cells were analyzed with the Double Color Enzymatic ELISPOT Assay according to the manufacturer’s instructions (ImmunoSpot). 3×10^5^ splenocytes per well were stimulated with Influenza HA peptide at 1 µg/mL (PM-INFA-HACal, JPT Peptide Technologies), 1 µg/mL ConA, or untreated in duplicate. Plates were imaged on the CTL Analyzer (Series 6 Universal) and processed with the ImmunoSpot software (Version 7.0.34.0). Counts per well were normalized to counts per 10^6^ splenocytes and background subtracted (untreated groups).

### HAI assay

Treated serum with RDE (Accurate Chemical, YCC340122) in saline (Moltox, 51-40S022.052) for 37°C for 16-22 hr. RDE was then inactivated by incubating samples at 56°C in a water bath for 30 min followed by cooling to RT for 15-30 min. Samples were diluted in PBS for a final serum dilution of 1:10. Whole turkey RBC (TRBC) (Lampire, 7209403) was rinsed in ice cold PBS and resuspended to 0.5% in PBS. Samples were serially diluted in PBS and 25 µL of sample mixed with 25 µL of 8 HAU virus (A/California/07/2009) and incubated at RT for 1hr. Added 50 µL of 0.5% TRBC and incubated at RT for 30 min. HAI titers were calculated as the reciprocal of the highest dilution resulting in 100% inhibition of RBC agglutination compared to the RBC only control. Samples were tested in 2-4 technical replicates. Values below the lower limit of quantification (LOQ) were transformed to 0.5*LOQ.

### Statistical analyses

Statistical tests were conducted in Prism (Version 10.4.0). Significance between groups from in vitro data was determined from a ratio paired t-test (p<0.05) and for in vivo generated data from Kruskal-Wallis test with Dunn’s multiple comparisons (p<0.05) as indicated in each figure legend.

## AUTHOR CONTRIBUTIONS

J.A.W, I.M., P.V.M., B.M., K.R.B., and T.F. conceptualized the study. J.A.W, R.M.J., I.M., B.F. P.V.M., P.M., and B.M contributed to the development and generation of reagents and assays. J.A.W, R.M.J., I.M., B.F. P.V.M., P.M, and B.M performed in vitro experiments. J.A.W, R.M.J, and B.F. managed in vivo studies and characterized immunogenicity. All authors were involved in study design and analysis. J.A.W. wrote the manuscript. All authors provided feedback and contributed to manuscript preparation.

## ACKNOWLEDGEMENTS

We thank Craig Wilen and Jonathan Smith for expert technical advice throughout the study and thoughtful comments during the preparation of this manuscript. We thank Jane Srivastava and the Gladstone Institutes Flow Cytometry core for expert assistance and support. We also knowledge the following funding sources for use of Gladstone Institutes Flow Cytometry core machines: NIH S10 RR028962 and the James B. Pendleton Charitable Trust for use of the Attune.

