## Supplemental info for "Controlling reactogenicity while preserving immunogenicity from a self-amplifying RNA vaccine by modulating nucleocytoplasmic transport"

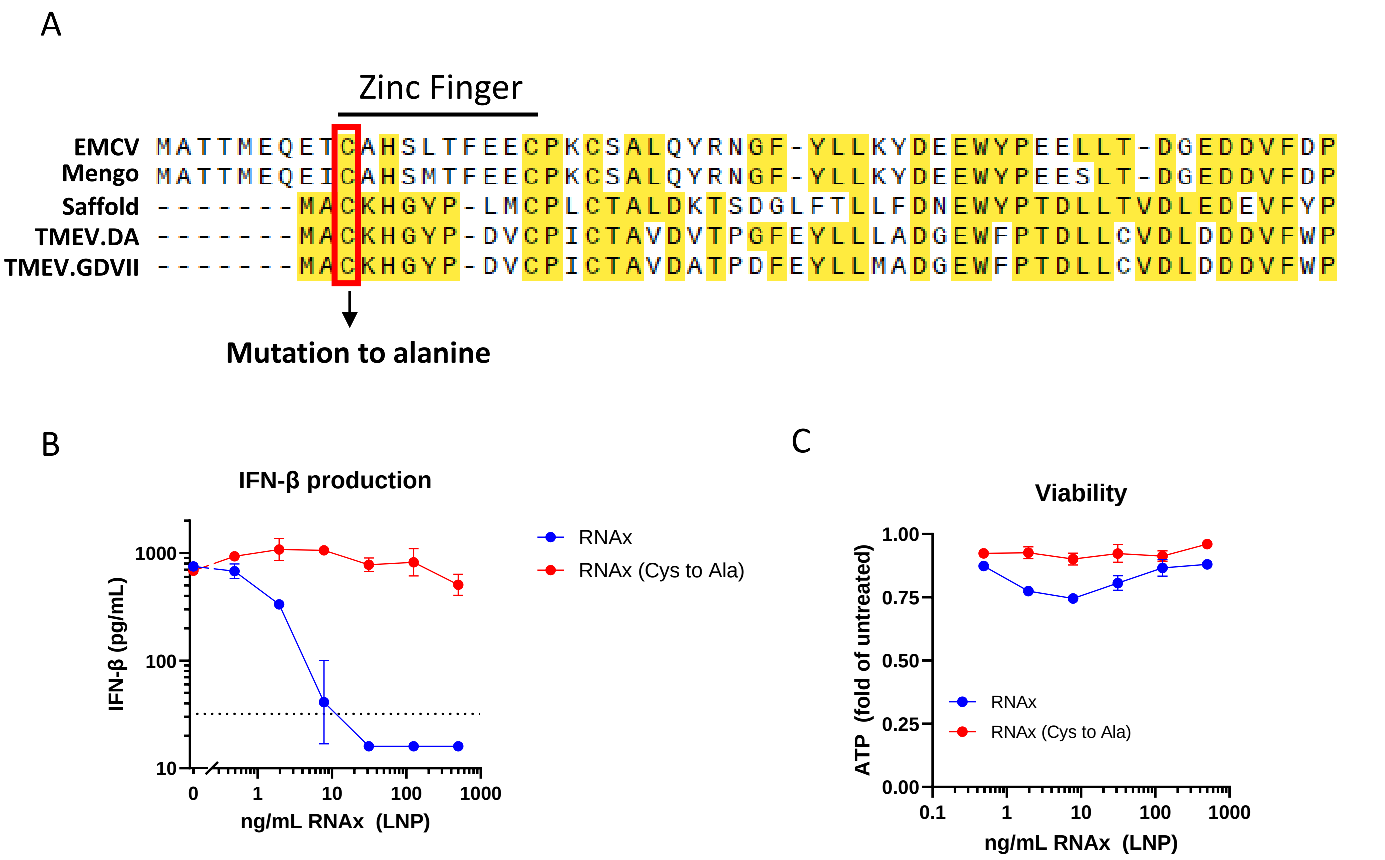


**Supplementary figure 1: Conserved Zinc finger domain of Cardiovirus leader protein is essential for suppressing IFN-β secretion via RNAx.** A) Amino acid alignment of leader proteins from major Cardioviruses EMCV (AHF20227.1), Mengo (AAA46547.1), Saffold (YP_001949875.1), TMEV.DA (AGM61326.1), TMEV.GDVII (NP_040350.1). Cys residue from the conserved zinc finger domain was mutated to Ala, then mRNA formulated into LNPs and added to BJ cells at various concentration for 24 hr in the presence of HA-nLuc saRNA-LNPs. B) IFN-β was quantified by Lumit assay (Promega) and C) viability by cell titer glo (Promega). Mean ± SEM, n=2

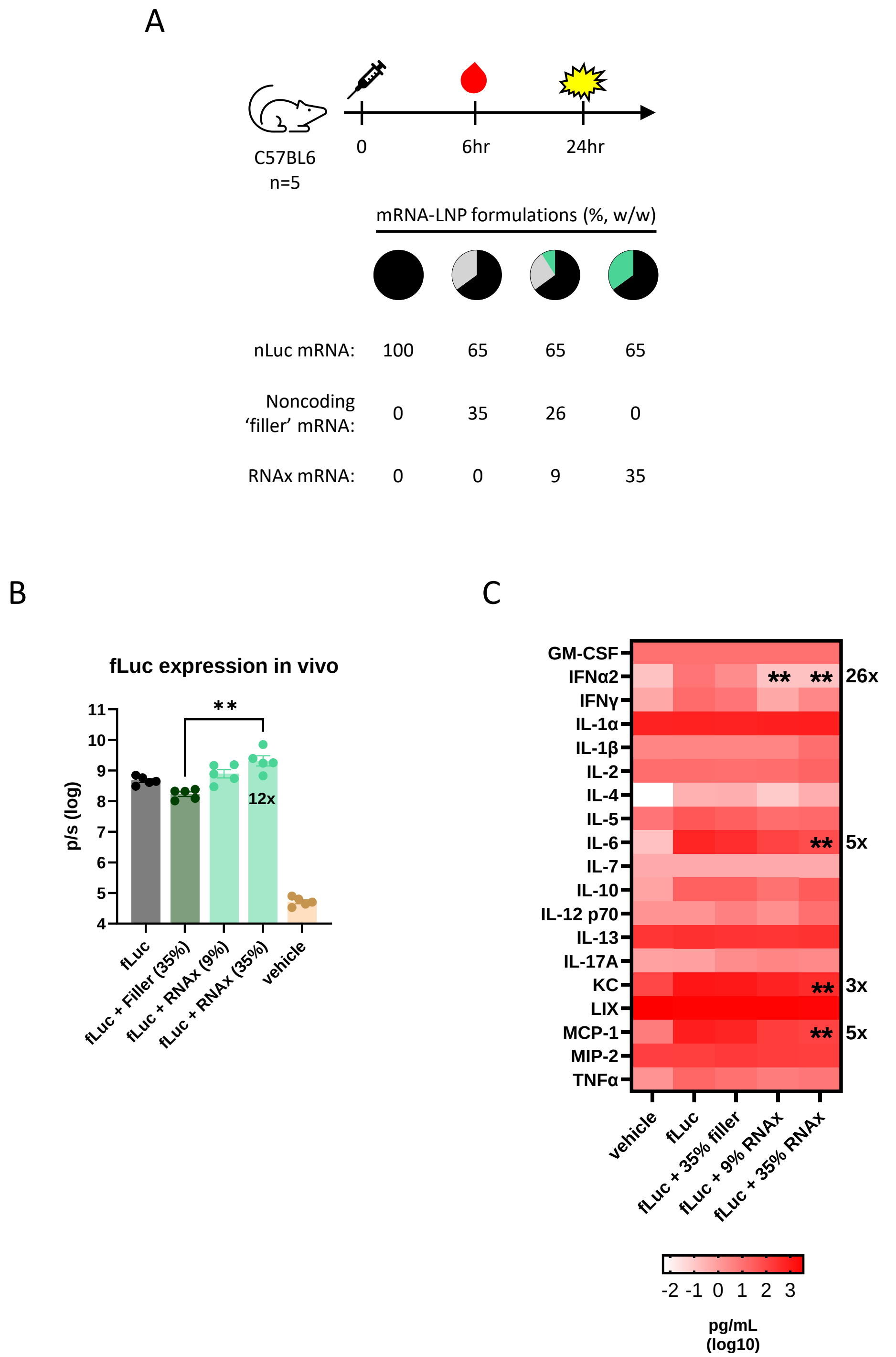


**Supplementary figure 2. RNAx enhances GOI expression and suppresses proinflammatory cytokines from unmodified RNA *in vivo.*** A) Unmodified fLuc mRNA was formulated into LNPs with or without doses of RNAx or a size-matched non-coding mRNA ('filler'). 5 µg mRNA-LNP was injected intramuscularly into C57BL/6 mice and 6hr later, serum was drawn for quantifying proinflammatory cytokines and 24 hr later, mice were imaged for Luc activity measured at the injection site with IVIS. B) Total flux of Luc signal is represented by photons per sec (p/s). Geometric mean ± geometric SD, n=5 mice per group, **p<0.01 Kruskal-Wallis test with Dunn's multiple comparisons (RNAx groups compared to fLuc + filler. Numbers on bar indicate average fold change enhancement of RNAx vs. filler. C) Serum cytokine levels quantified with a multiplex assay and ELISA (IFN-α2). Geometric mean. Asterisks indicate statistically significant reductions in cytokine levels (**p<0.01, Kruskal-Wallis test with Dunn's multiple comparisons) with the magnitude of the fold inhibition for each cytokine indicated to the right of the histogram.

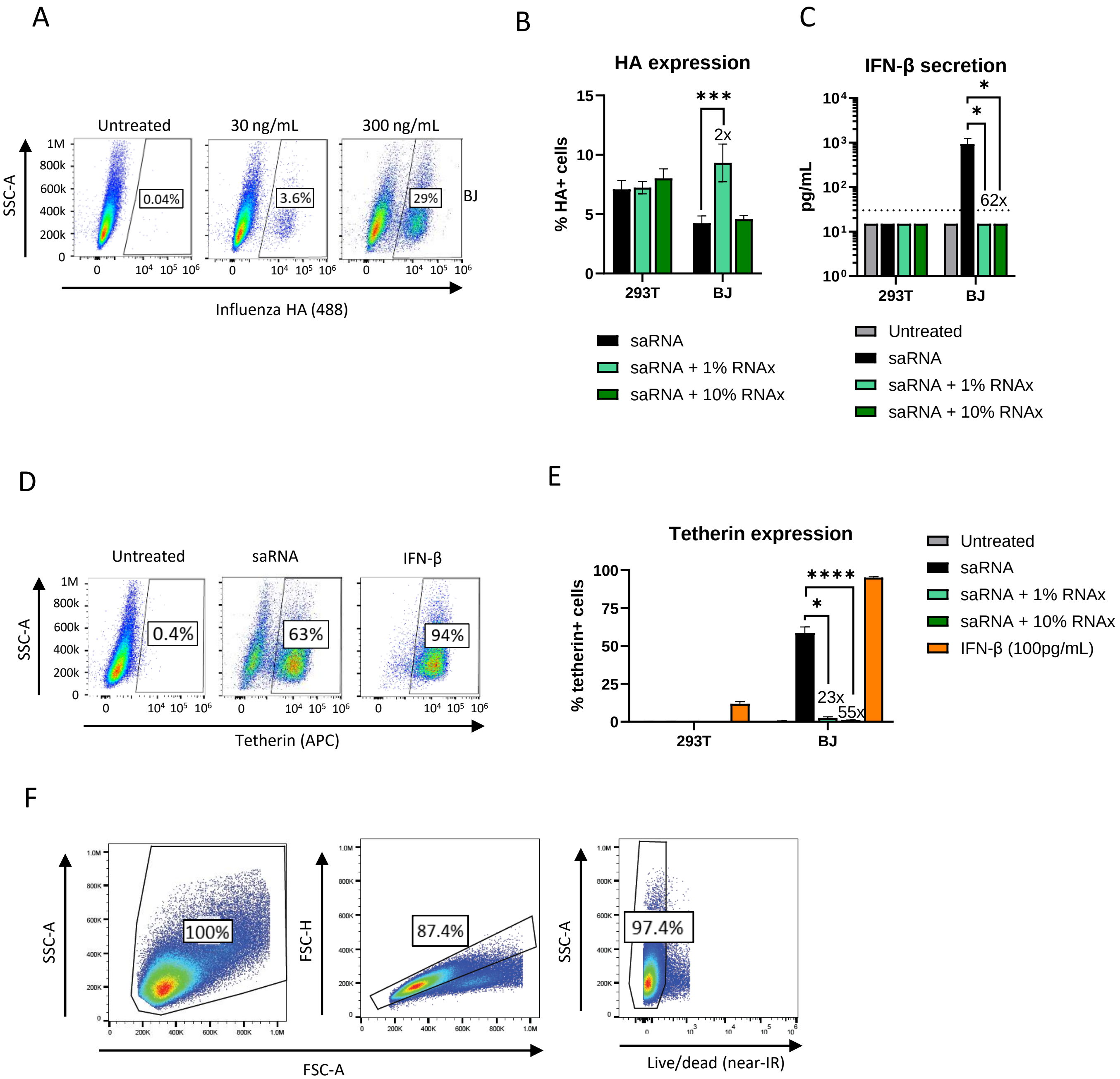


**Supplementary figure 3: *In vitro* expression of Influenza HA, tetherin, and IFN-β from HA saRNA-LNPs.** A) Representative flow cytometry plots for Influenza HA surface expression from HA saRNA-LNP treated BJ cells (no RNAx, 24 hr post treatment). B) 293T and BJ cells were treated with 30 ng/mL saRNA-LNPs with or with co-formulation with RNAx for 24 hr and Influenza HA surface expression quantified by flow cytometry and C) IFN-β secretion was quantified from the supernatants with ELISA. Dotted line = lower limit of quantification. D) Representative flow plots of surface tetherin staining by flow cytometry in BJ cell treated with 300 ng/mL HA saRNA-LNPs for 24 hr. 100 pg/mL of IFN-β served as a positive control. E) BJ cells were treated with 300 ng/mL saRNA-LNPs with or with co-formulation with RNAx for 24 hr and tetherin surface expression quantified by flow cytometry. Mean ± SEM, n=3, *p<0.05, ***p<0.001, ****p<0.0001 Ratio paired t-test. F) Representative plots of gating strategy for 293T (shown) and BJ cells. Numbers above bars indicate average fold change enhancement or inhibition of saRNA vs. saRNA.

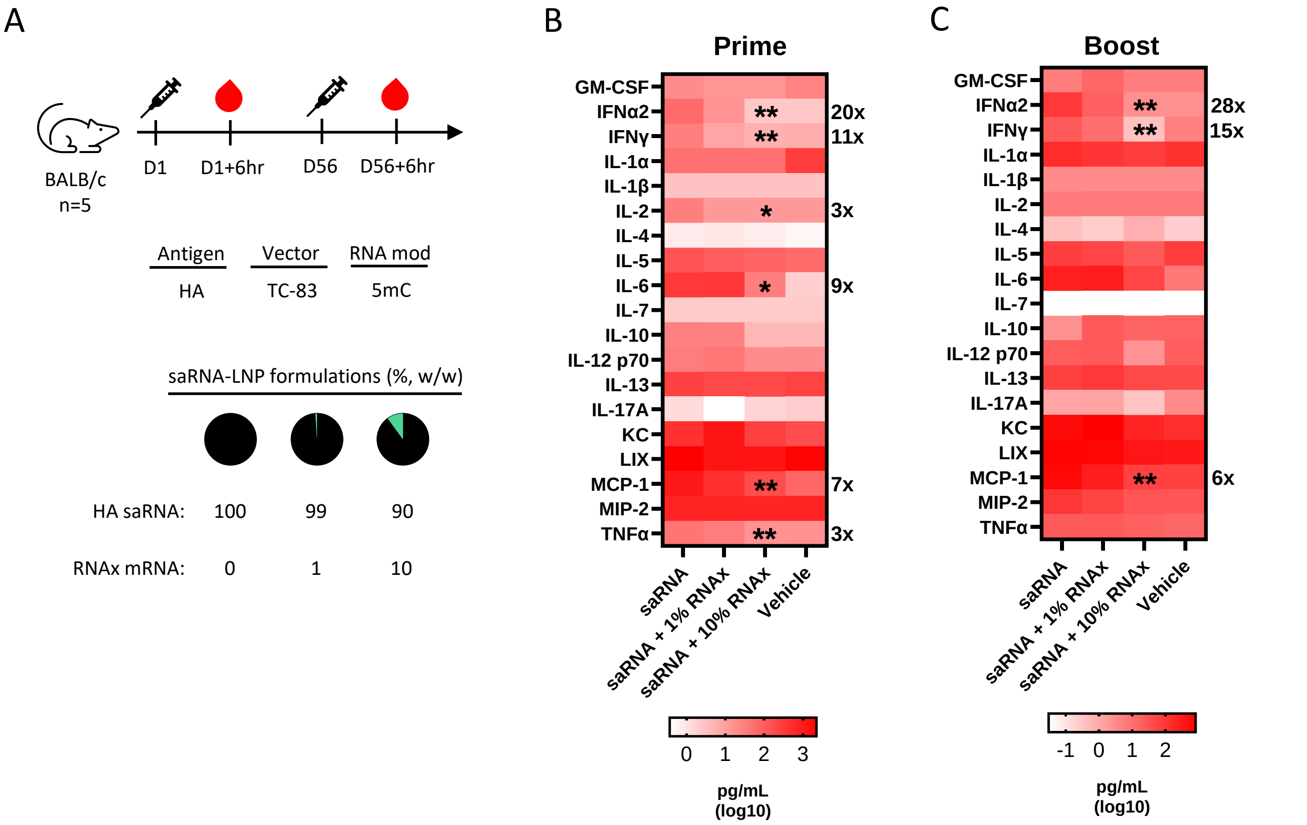


**Supplementary figure 4: RNAx suppresses proinflammatory cytokines in BALB/c mice following vaccination.** A) BALB/c mice (n=5 per group) were intramuscularly injected with 0.5 µg of Influenza HA saRNA-LNPs and were analyzed for serum cytokines 6 hr post injection following B) the prime or C) the boost at the indicated days of the study. Geometric mean, asterisks indicate statistically significant reductions in cytokine levels (*$p<0.05$, **$p<0.01$, Kruskal-Wallis test with Dunn's multiple comparisons) with the magnitude of the fold inhibition indicated to the right of each histogram.

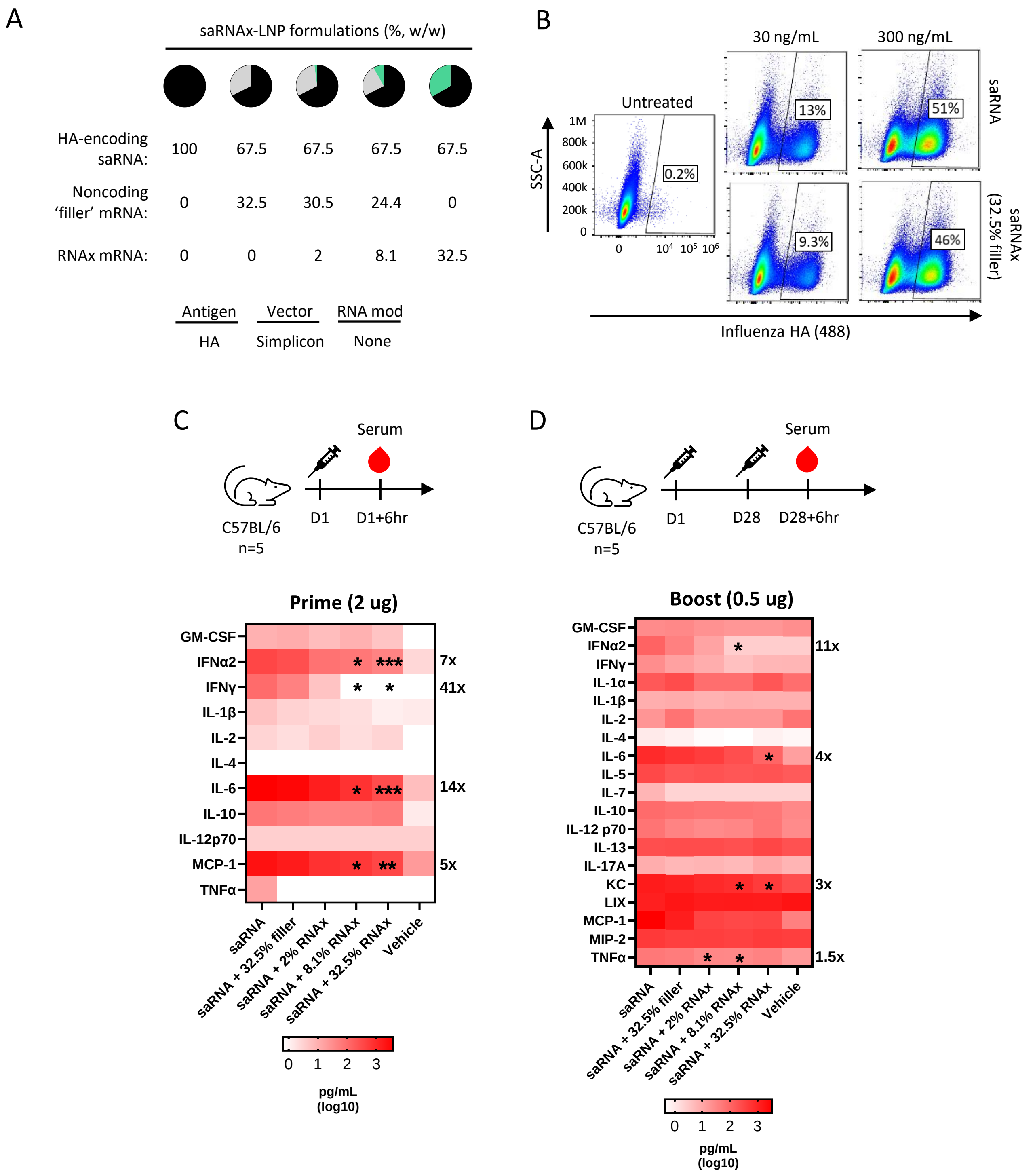


**Supplementary figure 5: RNAx suppresses proinflammatory cytokines in mice following vaccination with Simplicon saRNA vector.** A) Schematic of formulations. 'Filler' mRNA is a non-coding, sized matched (to RNAx) mRNA used to control for potential changes when formulating differently sized RNAs. Statistical tests were conducted vs. the 32.5% filler group. B) Flow cytometry of Influenza HA surface expression in 293T cells treated for 24 hr with HA saRNA-LNPs (no RNAx) from the Simplicon vector at the indicated concentrations. C57BL/6 mice (n=5 per group) were intramuscularly injected with Influenza HA expressing saRNA-LNPs and were analyzed for serum cytokines 6 hr post injection following A) the prime or B) the boost at the indicated days of the studies. Measurements for the prime and boost were generated from independent studies from incidental lack of sample collections, but from the same lot of LNPs. Animals from the prime samples were injected with 2 µg of saRNA-LNP while for the study where boost samples are analyzed, animals were primed and boosted with 0.5 µg of saRNA-LNP. Geometric mean, Asterisks indicate significant reductions in cytokine levels (*p<0.05, **p<0.01, ***p<0.001, Kruskal-Wallis test with Dunn's multiple comparisons) with the magnitude of the fold inhibition indicated to the right of each histogram.

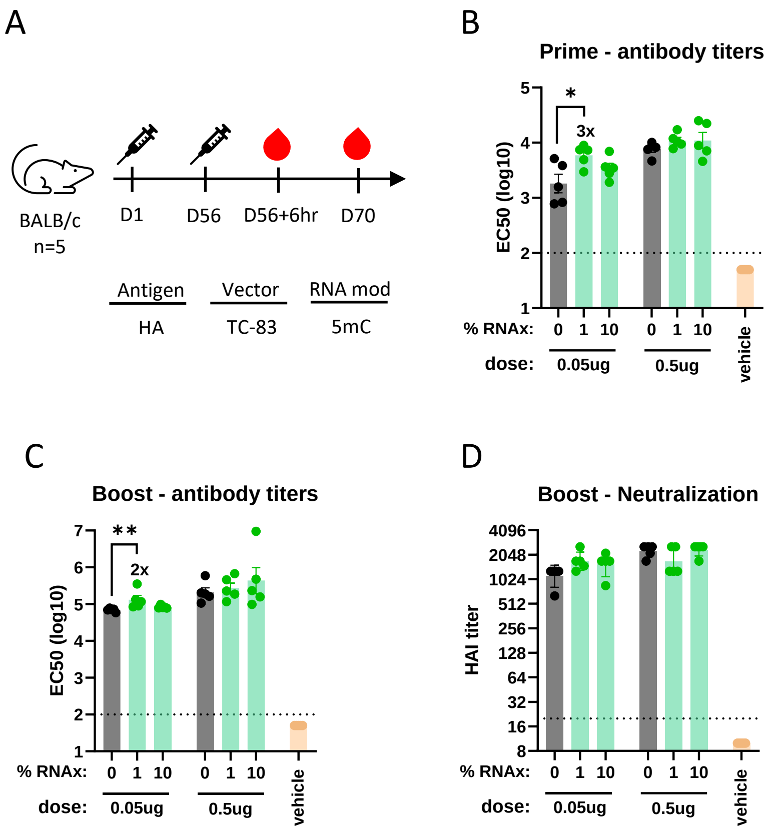


**Supplementary figure 6. Impact of RNAx on immunogenicity from Influenza HA saRNA-LNP vaccine in BALB/c mice.** A) BALB/c mice (n=5 per group) were intramuscularly injected (prime and boost) with 0.05 µg or 0.5 µg of HA saRNA-LNPs. Serum was collected 6 hr after the boost (post prime measurement) and serum and splenocytes collected 2 weeks post boost at the indicated days of the study. B) Total anti-HA IgG in the serum (EC50) was quantified by ELISA at post prime and C) 2 weeks post boost. D) Neutralization titers were quantified by the hemagglutination inhibition (HAI) assay 2 weeks post boost. Dotted line = lower limit of quantification, Geometric mean ± Geometric SD, $*p<0.05$, $**p<0.01$, Kruskal-Wallis test with Dunn's multiple comparisons. Numbers indicate fold enhancement of RNAx groups relative to saRNA only.

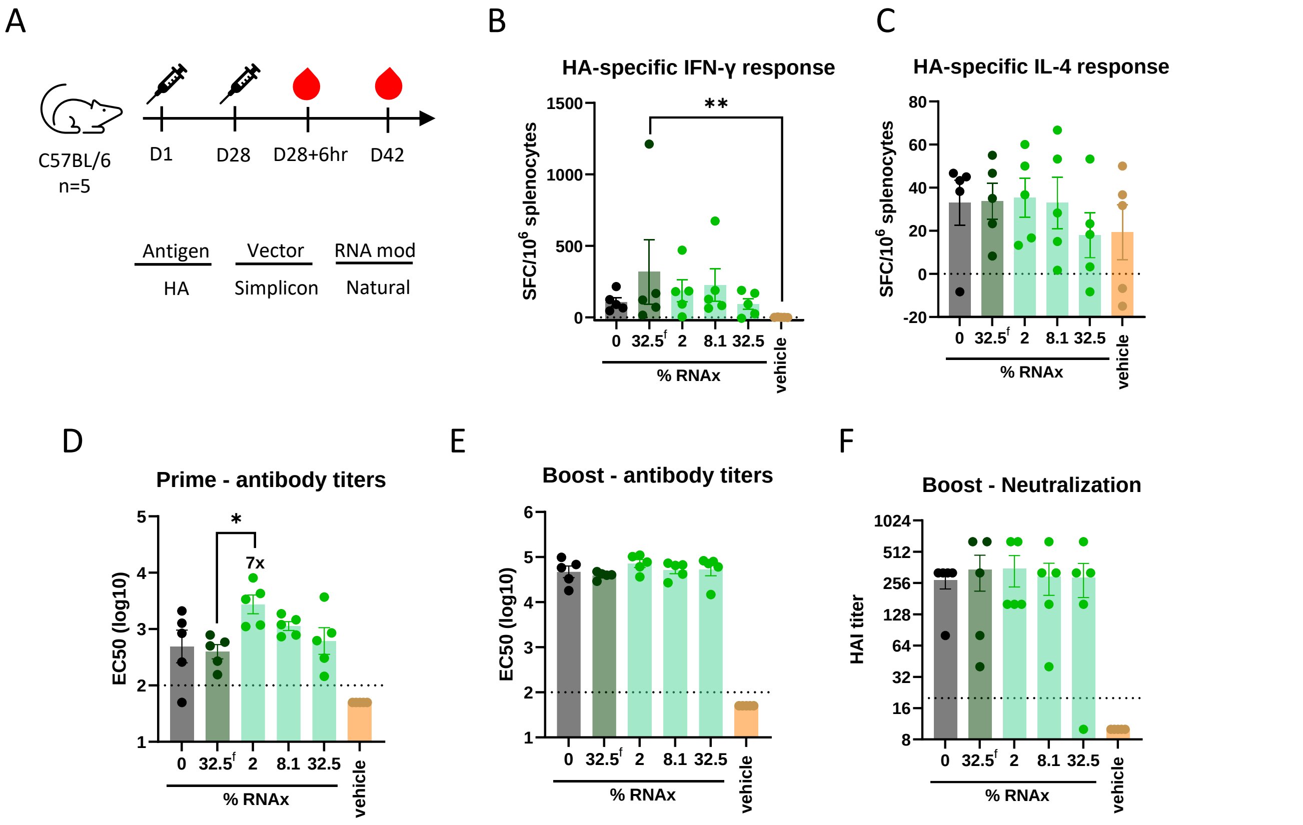


**Supplementary figure 7. Impact of RNAx on immunogenicity following vaccination with Simplicon saRNA vector.** A) C57BL/6 mice (n=5 per group) were intramuscularly injected (prime and boost) with 0.5 µg of HA saRNA-LNPs from the Simplicon vector (See Supplemental Fig. 4A). Statistical tests were conducted vs. the 32.5% filler group, denoted by $32.5^{f}$. Serum was collected 6 hr after the boost (post prime measurement) and serum and splenocytes collected 2 weeks post boost at the indicated days of the study. Splenocytes were treated with 1 µg/mL Influenza HA peptide pools and analyzed for B) IFN-γ and C) IL-4 secreting cells by ELISPOT. Spot forming cells (SFC) were subtracted by untreated samples and normalized by $10^6$ splenocytes. Mean ± SEM. D) Total anti-HA IgG in the serum (EC50) was quantified by ELISA at post prime and E) 2 weeks post boost. F) Neutralization titers were quantified by the hemagglutination inhibition (HAI) assay 2 weeks post boost. Dotted line = lower limit of quantification, Geometric mean ± Geometric SD, *$p<0.05$, Kruskal-Wallis test with Dunn's multiple comparisons. Numbers indicate fold enhancement of RNAx groups relative to saRNA only.

| Study | Treatment | size (nm) | PDI | EE% |
|---|---|---|---|---|
| HA-nLuc saRNA IVIS, PBMC | saRNA only | 128 | 0.14 | 86.3 |
| HA-nLuc saRNA IVIS, PBMC | saRNAx (36%) | 130 | 0.21 | 85.7 |
| Unmodified fLuc mRNA IVIS | fLuc | 94 | 0.14 | 82.3 |
| Unmodified fLuc mRNA IVIS | fLuc + Filler (35%) | 86 | 0.08 | 85.2 |
| Unmodified fLuc mRNA IVIS | fLuc + RNAx (9%) | 86 | 0.09 | 84.5 |
| Unmodified fLuc mRNA IVIS | fLuc + RNAx (35%) | 88 | 0.11 | 83.9 |
| HA saRNA (TC-83) immunogenicity and serum cytokines (C57BL/6 and BALB/c) | saRNA only | 148 | 0.17 | 82.8 |
| HA saRNA (TC-83) immunogenicity and serum cytokines (C57BL/6 and BALB/c) | saRNAx (1%) | 148 | 0.14 | 82 |
| HA saRNA (TC-83) immunogenicity and serum cytokines (C57BL/6 and BALB/c) | saRNAx (10%) | 157 | 0.14 | 80.8 |
| HA saRNA (Simplicon) immunogenicity and serum cytokines | saRNA only | 120 | 0.08 | 77 |
| HA saRNA (Simplicon) immunogenicity and serum cytokines | saRNA (filler, 32.5%) | 120 | 0.1 | 81.4 |
| HA saRNA (Simplicon) immunogenicity and serum cytokines | saRNAx (2%) | 120 | 0.11 | 84.1 |
| HA saRNA (Simplicon) immunogenicity and serum cytokines | saRNAx (8.1%) | 125 | 0.1 | 82.4 |
| HA saRNA (Simplicon) immunogenicity and serum cytokines | saRNAx (32.5%) | 119 | 0.12 | 82.2 |

**Supplementary table 1. Summary of RNA-LNP QC measurements.** Results from DLS, including average size, polydispersity index (PDI), and encapsulation efficiency (EE) percentage.

| Donor | Gender | Age | Race/ethnicity | Smoker | HBsAg | Anti-HCV | HIV-1,2 |
|---|---|---|---|---|---|---|---|
| 1 | Female | 25 | Caucasian | No | Neg | Neg | Neg |
| 2 | Female | 42 | African American | No | Neg | Neg | Neg |
| 3 | Male | 53 | Caucasian | No | Neg | Neg | Neg |
| 4 | Female | 34 | Mixed | No | Neg | Neg | Neg |

**Supplementary table 2. Demographics and health state of human PBMC donors obtained from commercial source.** PBMC cells from de-identified human donors were obtained from IQ Biosciences (cat# IQB-PBMC102) (Alameda, CA) and tested for Hepatitis B virus surface antigen (HBsAg), Hepatitis C virus antibody (anti-HCV) and HIV-1,2 by IQ Biosciences. Neg=negative.
